# Comparative genomics of *Exiguobacterium* reveals what makes a cosmopolitan bacterium

**DOI:** 10.1101/2020.11.19.390823

**Authors:** De-Chao Zhang, Zhaolu Zhu, Xudong Li, Ziyu Guan, Jinshui Zheng

## Abstract

Although the adaptation strategies of bacteria to specific environmental conditions are widely reported, fewer studies have addressed how microbe with cosmopolitan distribution adapted to diverse habitats. *Exiguobacterium* is a versatile genus whose members have been commonly found in great variety of habitats. To understand the mechanism behind the universally of *Exiguobacterium,* we isolated 103 strains from diverse environments, and performed large-scale metabolic and adaptive ability tests. We found that the capacities of survival in a wide range of temperature, salinity and pH are common for most *Exiguobacterium* members. According to the core genome based phylogeny and ANI analysis, 26 putative species including 13 putative new ones were identified and two genetic groups were classified as Group I and II. Comparative genomic analysis revealed that *Exiguobacterium* members can not only utilize a variety of complex polysaccharides and proteins that are ubiquitous in both terrestrial and marine environments, but also have a number of chaperonins and transporters which could support them to survive in different extreme environments. In addition, we found that the species from Group I can be found in more diverse environments with larger genome size compared to those of Group II. Twenty-five transporter families involved in transport of organic or inorganic substrates and environments stresses resistance were predicted to be enriched in Group I strains. This study provided the comprehensive insight into general genetic basis of the cosmopolitan distribution of a bacteria genus and deciphered putative determinants behind the ecological difference of different groups belonging to the same genus.

**IMPORTANCE:** The wide distribution characteristics make *Exiguobacterium* a valuable model for studying adaptive strategy of bacteria adapted to multiple habitats. In this study, we found that comprehensive capacity of diverse polysaccharides utilization and environmental stress resistance is the important basis for survival, and selective expansion of transporters is an evolution and adaptation strategy for extensive distribution. Our findings are significant for understanding the adaptation and evolution mechanisms of cosmopolitan bacteria and explaining the vital genomic traits that facilitate niches adaptation.

## INTRODUCTION

Microbial community composition and diversity across landscape is nonrandom (1). Physical and chemical factors in the environment significantly influence the distribution patterns of microbes (2). There is a barrier between marine and non-marine habitats due to their strong physiochemical difference such as salinity, temperature, pH, dissolved oxygen and water chemistry (3). As a result, most marine microbes belong to different phylogenetic groups from their freshwater and terrestrial relatives, and there are rare transitions between these two niches (4, 5). It was frequently reported that these different bacteria members usually utilize different strategy for niche adaptations. Comparative genomics of ocean microbes revealed that many marine bacteria were identified with streamlined genome to reduce metabolic costs of maintaining nonessential genetic material to adapt to the nutrient-poor ocean environments (6, 7). And it is obviously different from free-living terrestrial bacteria, which usually have normal genome size with frequent horizontal genetic transfer events to facilitate the capacity of using diverse collection of nutrients and ability of resistance to complicated adverse environments (8, 9). The transition events of marine to non-marine or in reverse for bacteria require complex genome evolution (10, 11). Events of gaining and losing gene which involved in the abilities for transport, metabolism and assimilation different types of organic or inorganic nutrients play crucial roles during this progress (10, 12). However, this knowledge is mainly derived from the comparative genomics of a few bacteria with high abundance in either marine or non-marine microbiota (13, 14). The strategies of evolution and adaptation for microbes which have wide distribution in both marine and non-marine environments are not well studied yet.

Bacteria of the genus *Exiguobacterium* are Gram-positive facultative anaerobes which have been frequently isolated from various habitats including seawater, marine sediment, marine algae (15–17), soil (17), freshwater (18), plant rhizosphere (19), and some extreme environments such as salt lake (20), glacier and hot spring (21). Genomic analysis of these bacteria has provided some vital insights into their psychrophilic and thermophilic adaptations and multiple toxic compound resistances (22–24). However, comprehensive knowledge about these bacteria evolutionary adaptation to marine and non-marine habitats remains undiscovered.

In this study, *Exiguobacterium* was used as a model to study how microbes with cosmopolitan distribution adapted to both marine and terrestrial environments. We leveraged different strategies to decipher the ecology and evolution of *Exiguobacterium* spp.. We firstly mined the public database for 16S rRNA gene to reveal the diversity and distribution of the genus. Then we isolated multiple strains from marine and other habitats and tested their adaptive and metabolic features. Furthermore, we sequenced 103 strains and performed large-scale phylogenomic and comparative genomic analyses on a total 145 genomes representing strains isolated from marine and non-marine habitats worldwide. Special attention was drawn on genomic and metabolic characters responded to diverse environments.

## RESULTS AND DISCUSSION

### Cosmopolitan distribution of *Exiguobacterium* spp. and their survival abilities in a wide range of conditions

To explore the diversity and distribution of members belonging to *Exiguobacterium*, we focused on the bacterial strain with 16S rRNA gene sequence having an identity above 95% to that of the reported species type strain from this genus. A total of 2,582 *Exiguobacterium* 16S rRNA gene sequences with unambiguous information of isolation source were collected from Genbank (Table S1). We found that members of this genus were frequently isolated from territorial environments (86.6%), including plant or rhizosphere (16.6%), animal skin or gut (10.7%), freshwater or freshwater sediment (12.7%), contaminated water or soil (7%), soil (6.8%), extreme environments (hot- or cold-associated or hypersaline environments) (6%), air (4.8%) and other non-marine environments (22%) (Fig. 1A; Table S1). The left 13.4% members were isolated from marine-associated environments including sea water, algae and oceanic sediment. Combining the location information, we found that *Exiguobacterium* can be isolated from all of the continents and oceans. This result was accordance with the current notion that *Exiguobacterium* was a cosmopolitan bacteria genus including many extremophiles surviving in both marine and non-marine environments worldwide (15).

**Fig. 1.**
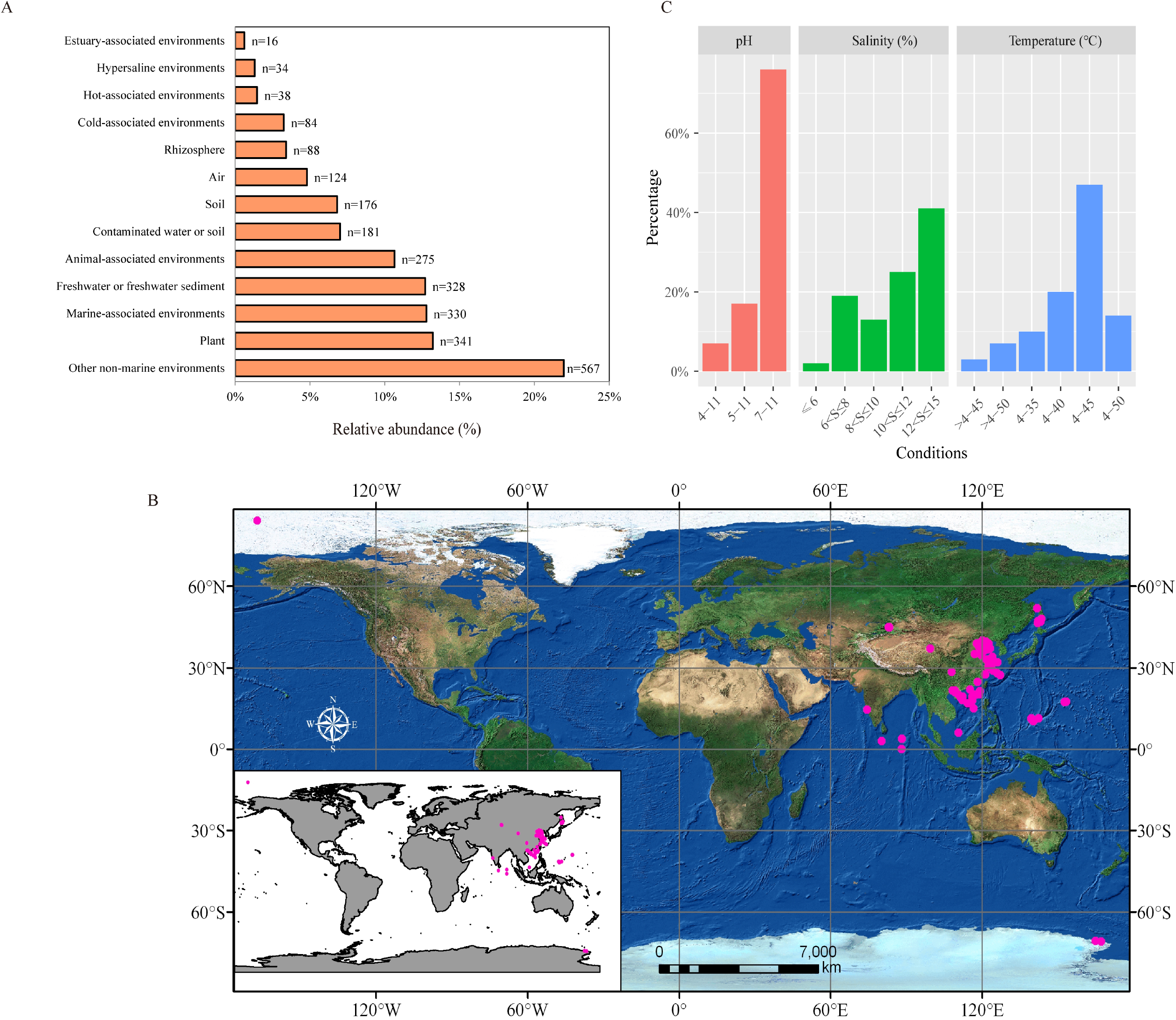
Cosmopolitan distribution of *Exiguobacterium* strains. (A) Relative abundance of 16S rRNA gene sequences among 13 types of habitats. (B) The isolation sites of 98 *Exiguobacterium* strains in this study (The pink dots represent the sampling location). (C) Temperature, pH and salinity tolerance test of 103 *Exiguobacterium* strains. In the temperature experiment, the intermediate temperature between 4°C and 25°C was not tested. The lowest growth temperature was not recorded for the strain that could not grow at 4°C.

To study the adaptation and evolution of *Exiguobacterium,* we performed large-scale phenotype tests and comparative genomics analysis. A total of 103 *Exiguobacterium* strains were collected here, including 87 isolates from marine-associated niches (marine sediment, seawater, algae, marine cold spring, hydrothermal vent, seamount, mangrove, marine fish and coral), 11 strains from territorial environments (soil, salt lake, coal mine and pig farm) and 5 type strains (Fig. 1B; Table S2). Among the 42 *Exiguobacterium* members with genome available up to the manuscript submitted, only 4 strains were isolated from marine habitats. We used more isolates from marine than territorial environments in this study.

Previous studies have reported that *Exiguobacterium* spp. could survive in a wide range of habitats including cold, hot, hypersaline and alkaline environments (16), while it is unclear if these features are shared by all the members or strain/species specific.

To assess the survival abilities of *Exiguobacterium* spp. to diverse conditions, we evaluated the growth potential of the collected 103 strains in culture medias with different of pH, temperatures and salinity, respectively. We found that the most *Exiguobacterium* members can survive and grow in a wide range of temperature, salinity and pH values, respectively. (Fig. 1C; Table S3). The pH test revealed that all the strains were alkali-resistant which could survive in the environments with pH value up to 11, and 24% strains showed growth at pH 5 (17%) and even 4 (7%). For salinity test, most of strains showed tolerance to saline, with 66% strains growing in environment with NaCl concentration above 10%. Temperature tests showed that the *Exiguobacterium* spp. have high and low temperature tolerance, with 91% strains growing at *4°C* and 61% surviving from 40 to 50°C. Moreover, no associations were found between the growth abilities and the source environments. These results suggest the extensive adaptability to survive in variable environments as a general feature of this genus.

### Phylogenetic analysis identified two genetic groups in *Exiguobacterium* genus

We conducted comparative genomics analysis to investigate the strategies of evolution and adaptation to diverse environments. All genomes of the 103 collected *Exiguobacterium* strains were sequenced through Illumina Novaseq 6000 platform and assembled by SPAdes software (25). The 42 *Exiguobacterium* genomes available on GenBank were also included in the comparative analysis (Table S2). All the annotated proteins from 145 genomes were clustered into 8,728 groups, with 1,162 shared by all genomes classified as core gene families. The maximum-likelihood (ML) phylogenetic tree was constructed based on the core genome alignment (Fig. 2). Two genetic groups of *Exiguobacterium* were classified and well-supported by bootstrapping analysis, which is consistent with previous analysis based on 16S rRNA gene sequences (15). Using the threshold of ANI (average nucleotide identity) 95% to define different species (26), a total of 26 species can be classified including 13 putative new ones (N1 to N13) (Fig. 2; Table S4). There were 11 and 15 species belonging to Group I and II, respectively (Table S4).

**Fig. 2.**
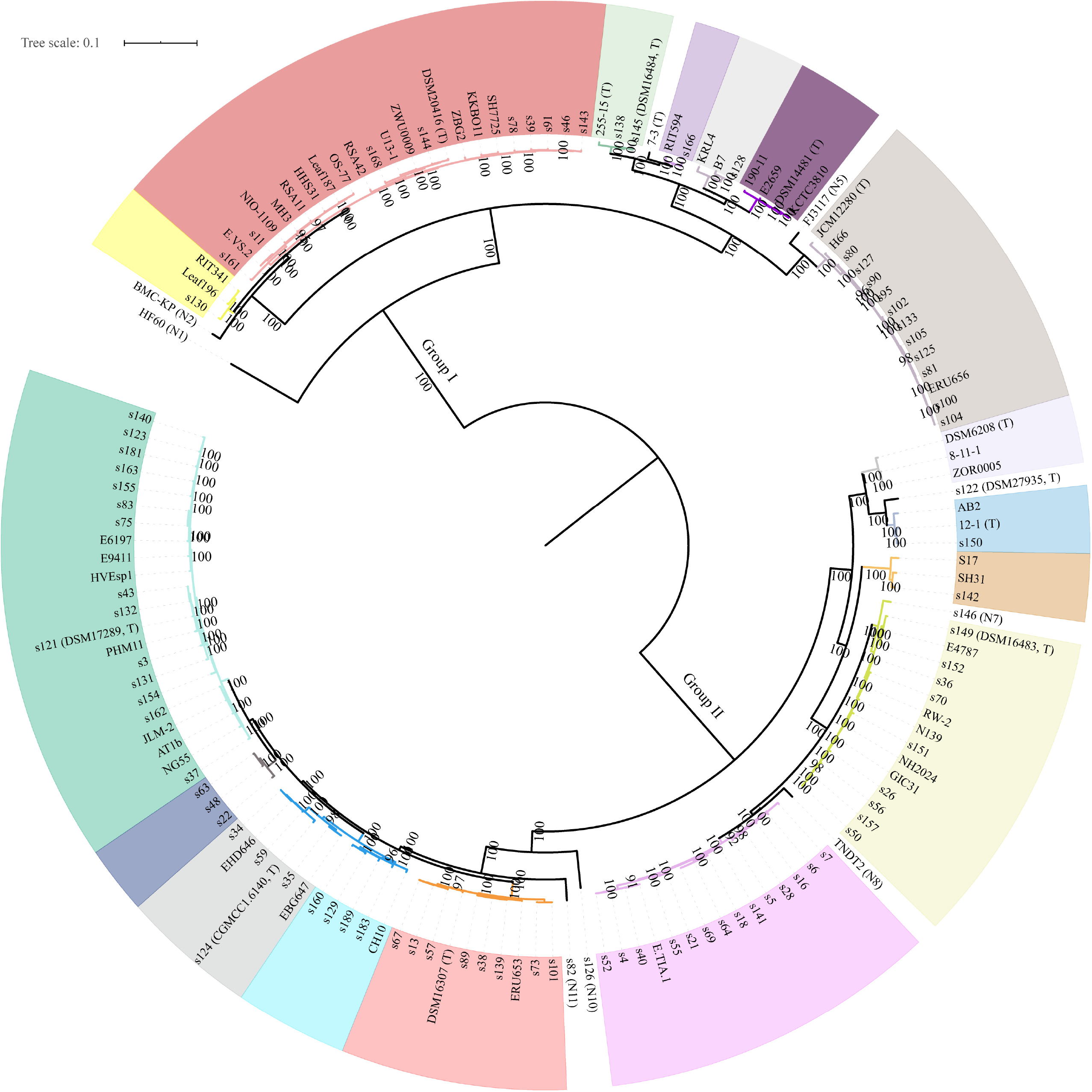
Phylogenetic analysis of *Exiguobacterium*. The tree was built using IQ-tree based on the concatenated amino acid sequence alignments of 1,162 core genes. Bootstrap support values were calculated from 1000 replicates. ‘T’ represents type strain: NIO-1109 for *E. enclense,* HHS31 for *E. indicum,* DSM20416 for *E. acetylicum*, 255-15 for *E. sibiricum*, s145 for *E. artemiae*, 7-3 for *E. sibiricum*, DSM14481 for *E. undae*, JCM12280 for *E. oxidotolerans*, DSM6208 for *E. aurantiacum*, s122 for *E. himgiriensis*, 12-1 for *E. alkaliphilum*, s149 for *E. mexicanum*, DSM16307 for *E. marinum*, s124 for *E. aestuarii* and s121 for *E. profundum*. N1 to N13 represents putative new species.

We next annotated the tree by adding the isolated environment of each strain, and found that strains from different marine and terrestrial niches distributed around the whole tree, even isolates from the same species could found in different environments (Table S5). For example, members of *E. acetylicum* were found in seawater, ocean sediment, soil, rhizosphere, glacier and even animal gut. It is suggested that frequent transitions *Exiguobacterium* spp. among different niches of terrestrial and marine environments. This finding is different from those of some typical marine bacteria which usually contained different lineages adapting to marine and non-marine habits, respectively (27).

### Carbon and nitrogen source utilization for wide adaptation

To explain the extensive distribution of *Exiguobacterium* strains in different niche types, we focused on the genes involved in different types of nutrition metabolism. Active enzymes involved in carbohydrate metabolism are defined as carbohydrate-active enzymes (CAZymes). A total of 7,864 genes belonging to 5 CAZyme superfamily were identified from all the genomes, with 61.5%, 18.7%, 15.9%, 2.7% and 1.2% genes belonging to GH (glycoside hydrolase), CE (carbohydrate esterase), CBM (carbohydrate-binding module), PL (polysaccharide lyase) and AA (auxiliary activities), respectively (Fig. 3A; Table S6). Each genome sequence encoded 43 to 68 of these enzymes.

**Fig. 3.**
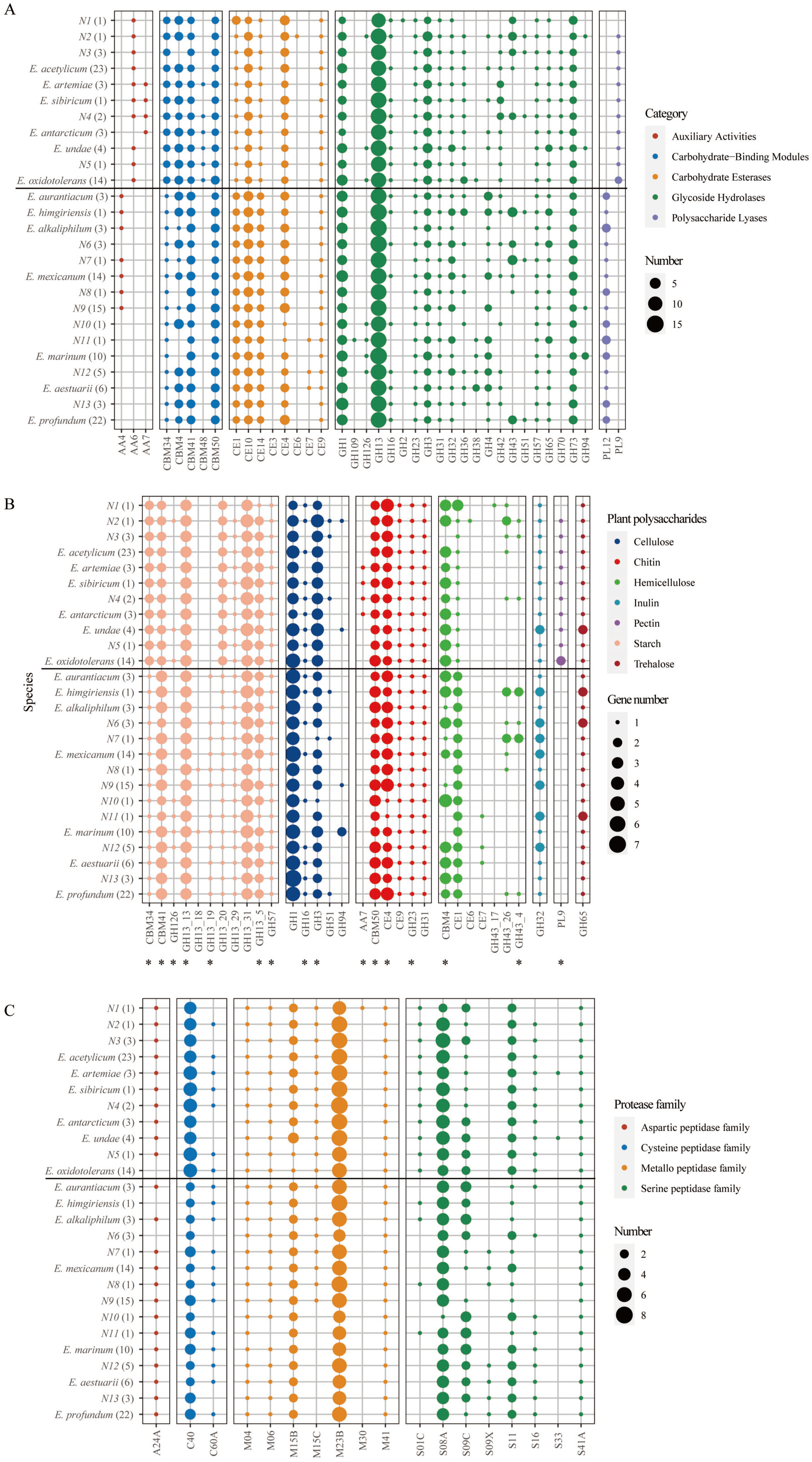
Carbon and nitrogen source utilization. (A) Number of carbohydrate-active enzymes (CAZymes) encoded in *Exiguobacterium* genomes. (B) Number of plant polysaccharides degradation enzymes encoded in *Exiguobacterium* genomes. (C) Number of extracellular peptidases encoded in *Exiguobacterium* genomes. Brackets: Total number of genomes in each *Exiguobacterium* species. Asterisk: CAZyme with potential secretion signal.

Many classes of enzyme for complex polysaccharide degradation were predicted for *Exiguobacterium* genomes (Fig. 3B; Table S6). The top 3 abundant classes are those associated with the degradation of starch, cellulose and chitin, and most of these CAZymes are potentially secreted (Fig. 3B). Family GH13 represents the main amylolytic enzymes family, the GH13_31 (α-glucosidase) and GH13_13 (pullulanase) are the top 2 frequent subfamilies, with 3.3 and 3 genes per genome, respectively. The most abundant family involved in cellulose degradation is GH1 (β-glucosidases), with each genome containing more than 4 genes. For chitin degradation, family CE4 (deacetylase) and CBM50 (chitin-binding) showed significantly higher abundance.

As the important storage polysaccharide, starch is produced by plants from both terrestrial and marine (28, 29). Cellulose is the most prevalent polysaccharide in nature, which make up the plant and algal cell walls (30, 31). As the second common polysaccharide after cellulose in nature, chitin is also widely distributed in terrestrial and marine ecosystems as a major structural component of crustacean shell, arthropod exoskeleton and the cell wall of diatom (32–34). The abundant CAZymes contained by members of *Exiguobacterium* giving the strong putative capacity for degradation of these polysaccharides ensure these bacteria to extensively gain the carbon source and survive in variable niches of both marine and non-marine environments.

Proteinaceous compounds are abundant forms of organic nitrogen in aquatic and soil (35). Extracellular microbial peptidases play an important role in both marine and terrestrial environments, as they directly link to organic nitrogen degeneration to contribute the global nitrogen cycling (35). In this study, a total of 3,912 putatively secreted peptidases were assigned to 20 families, including 43.7%, 38.1%, 15% and 3.2% belonging to metallo-, serine, cysteine and aspartic peptidase families, respectively (Fig. 3C; Table S7). When normalized to genome size, the average number of secreted peptidase coding genes was 9 genes per Mb, which is higher than the overall level of bacteria (5.84 genes per Mb) (35). Among these peptidases, the metallo peptidase M23 and serine peptidase S08 represent the top 2 ample peptidases (Fig. 3C). M23 peptidases were reported to degenerate the bacterial extracellular peptidoglycan, contributing to nutrition acquisition or defense against competitors (36, 37). Serine peptidases are often used as marker enzyme for proteolysis activity in soil, and play important roles in the utilization of nitrogen sources in the environments (38). The presence of abundant potentially secreted peptidases in *Exiguobacterium* genomes could allow them to exploit different niches for nitrogen source uptake in different environments.

To validate the potential ability of *Exiguobacterium* spp. to degrade and metabolize complex carbohydrates and proteins, the amylase and protease activities of the 103 strains isolated in this study were tested on plates (Fig. S1). All of these strains showed effective hydrolysis ability for starch, and approximately 70% of these strains can degenerate proteins. Taking results from both the genomic analysis and activity testing together, it provided strong evidence that most members of the *Exiguobacterium* genus have the abilities to metabolic and utilize a wide range of nutrition from marine and non-marine environments, which explains the genetic basis for the cosmopolitan distribution of these bacteria.

### Genetic basis of keeping homeostasis in extreme environments

As a versatile genus, *Exiguobacterium* was found to survive in many extreme environments such as cold, hot, saline and pollutant (16). Our phenotype test has proved that these adaptive characteristics are shared by most members of this genus. We investigated the putative genetic determinants behind these abilities from the whole genus.

Two strategies are used by bacteria to survive in cold environment, which are utilization of unsaturated branched-chain fatty acids to maintain membrane fluidity and expression of cold shock proteins (Csp) that stabilize the bacterial cytosol at low temperatures (39, 40). From the genomic analysis, we found that all members of *Exiguobacterium* genus could use the both strategies to cope with low temperature. Two types of fatty acid desaturase (FAD) involved in unsaturated branched-chain fatty acid production were identified in *Exiguobacterium* genomes (Fig. 4; Table S8). All genomes except those of AB2 and s126 encoded at least one FAD1 protein, while gene for FAD2 was mainly contained by the strains belonging to Group I. Three types of *csp (cspA, cspB* and *cspC)* were predicted from all the genomes (Fig. 4). Most genomes in Group I contained more than 2 *cspA* genes while those from Group II had only one. The *cspB* and *cspC* were harbored by members from Group I and II, respectively. It was reported that members can grow below 0°C mainly belonging to Group I (15). More *csp* and *fad* genes contained by member of Group I than II may contribute to this difference.

**Fig. 4.**
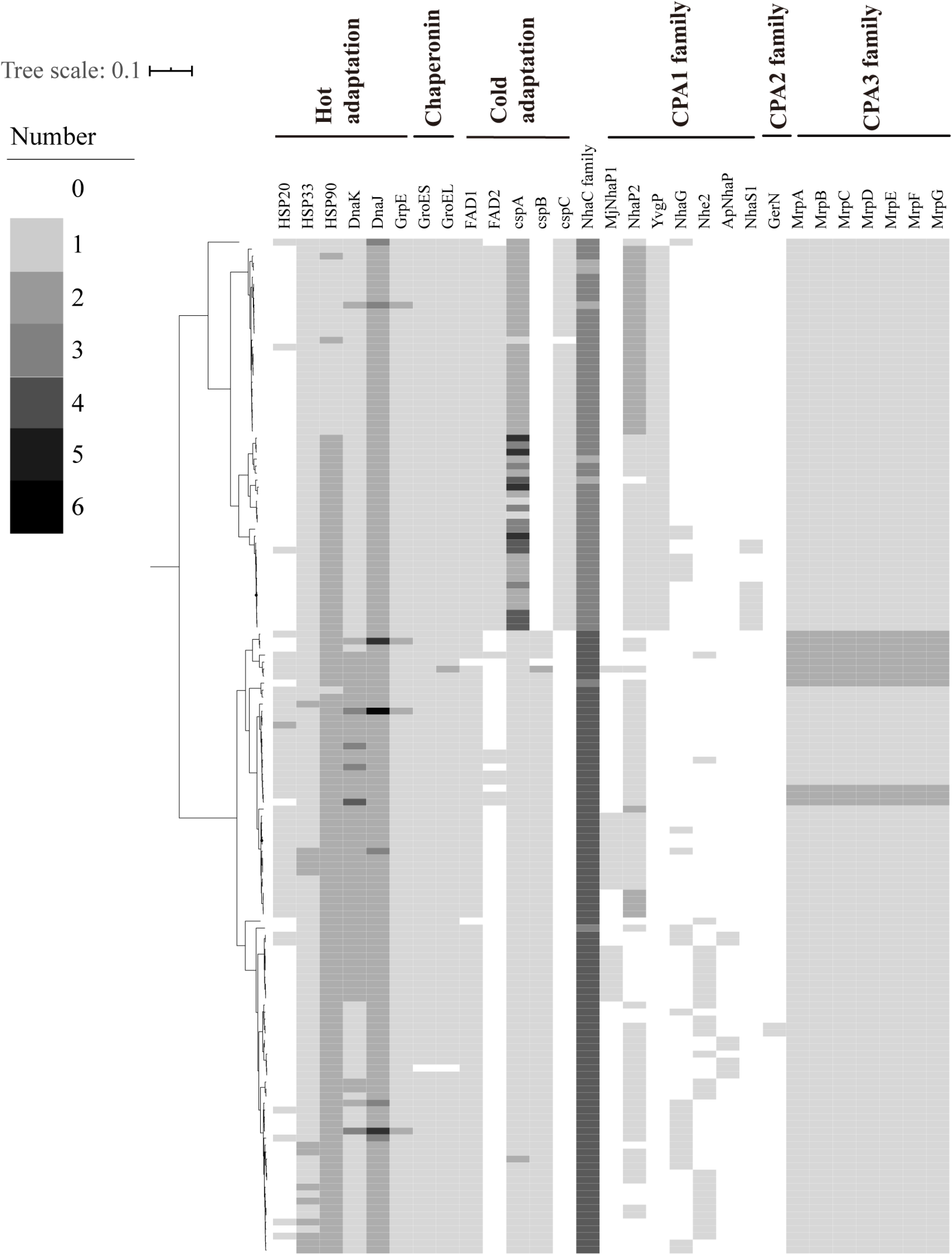
Vital genes detected across *Exiguobacterium* genomes for keeping homeostasis in extreme environments. The heatmap represents the vital gene number with distribution across the 145 genomes. The maximum-likelihood tree was constructed by RAxML as described in methods.

To support to survive in hot environments, all strains of *Exiguobacterium* spp. contain the shock gene cluster *grpE-dnaJ*-*dnaK* (Fig. 4; Table S8), which encodes chaperones that prevent aggregation and denaturation of proteins at high temperature (41). It was reported that proteins GroEL/GroES cooperating with DnaK/DnaJ to prevent protein misfolding in bacteria (42). Genes encoding GroEL and GroES were also discovered in all of the strains except for EHD646 (Fig. 4; Table S8). Additionally, other 3 types of heat shock protein (HSP20, HSP33 and HSP90) were also predicted. These HSPs are important chaperone for appropriate response to heat or oxidative stress because of their capacity of preventing irreversible protein denaturation (43–46). This suggests that members of this genus utilize multiple ways to cope with heat environments.

In bacteria, the Na^+^:H^+^ antiporters play crucial roles in the maintenance of intracellular pH homeostasis and dynamic balance of cellular Na^+^. According to the Transporter Classification Database (TCDB), Na^+^:H^+^ antiporters mainly contain the large monovalent cation/proton antiporter (CPA) family such as CPA1, CPA2 and CPA3, and the NhaC Na^+^:H^+^ antiporter family (47, 48). Seven types of CPA1 and 1 from CPA2 of the Na^+^:H^+^ antiporters of were predicted from the genomes of *Exiguobacterium* spp. (Fig. 4; Table S8). The Na^+^:H^+^ antiporters from CPA1 and CPA2 family were found in partial *Exiguobacterium* genomes, with the former one more frequently from ones of Group I and the last more common from those belonging to Group II. Compared to CPA1 and CPA2, CPA3 antiporters are more structurally complex with a multicomponent structure consisting of either seven or six members (49). This multicomponent Na^+^:H^+^ antiporter (Mrp) has been proved to provide Na^+^/H^+^ antiport activity and function in multiple compound resistance and pH homeostasis in *Bacillus subtilis* (49). In this study, Mrp antiporters were identified from all *Exiguobacterium* genomes (Fig. 4; Table S8). In addition, the antiporter from NhaC Na^+^:H^+^ antiporter (NhaC) family was identified in all *Exiguobacterium* genomes with copy number up to 6 (Fig. 4; Table S8). The presence of multiple types of Na^+^:H^+^ antiporter provides the basis for *Exiguobacterium* to maintain osmosis and pH balance in a variety of environments.

The combining results of the existence of diverse important proteins, including cold- and heat-shock protein, chaperonin, fatty acid desaturase and diverse Na^+^:H^+^ antiporters could explain the broad range of acceptable temperatures, pH and salinity for *Exiguobacterium* strains, and help to colonize in diverse habitats.

### Expansion of transporter families contributing to a wider adaptation

By comparing the ecological difference between members of Group I and II, we found that strains from marine environments were more frequently assigned to Group II and species from Group I had a more diverse niche distribution. Based on the analysis of 16S rRNA gene sequences from GenBank, there were 54 and 292 respectively belonging to Group I and II among the 346 sequences from marine environments (Table S1). As for the strains isolated in this study, there were 25 and 62 respectively belonging to Group I and II among the 87 strains from marine environments. In Group I, 10 of the 11 species contained strains isolated not only from the marine environment, but also from various terrestrial environments such as soil, plant rhizosphere, fresh water, etc (Table S5). While in Group II, most species were mainly isolated from marine-associated environments (Table S5). More diverse niche distribution suggested that the members belonging to Group I have a stronger capacity for environments adaptation.

To understand the genetic background behind the ecological difference of these two groups, we performed comparative genomic analysis and found a similar increasing tendency of genome size and transporter number from species of Group I to Group II (Fig. 5A; Fig. 5B and Table S9). The average genome size and transporter number of Group I (3.12Mb and 648) was significant larger than those of Group II (2.90Mb and 610) (*p* < 0.0001, Wilcoxon test) (Fig. 5C; Fig. 5D). Moreover, the spearman correlation coefficients of transporter number with genome size and CDS number is 0.86 (Fig. 5E). The significant correlation (*p* < 0.01) with genome size and CDS number suggests that the expansion of transporter contributed to the difference of genome content in Group I and Group II strains.

**Fig. 5.**
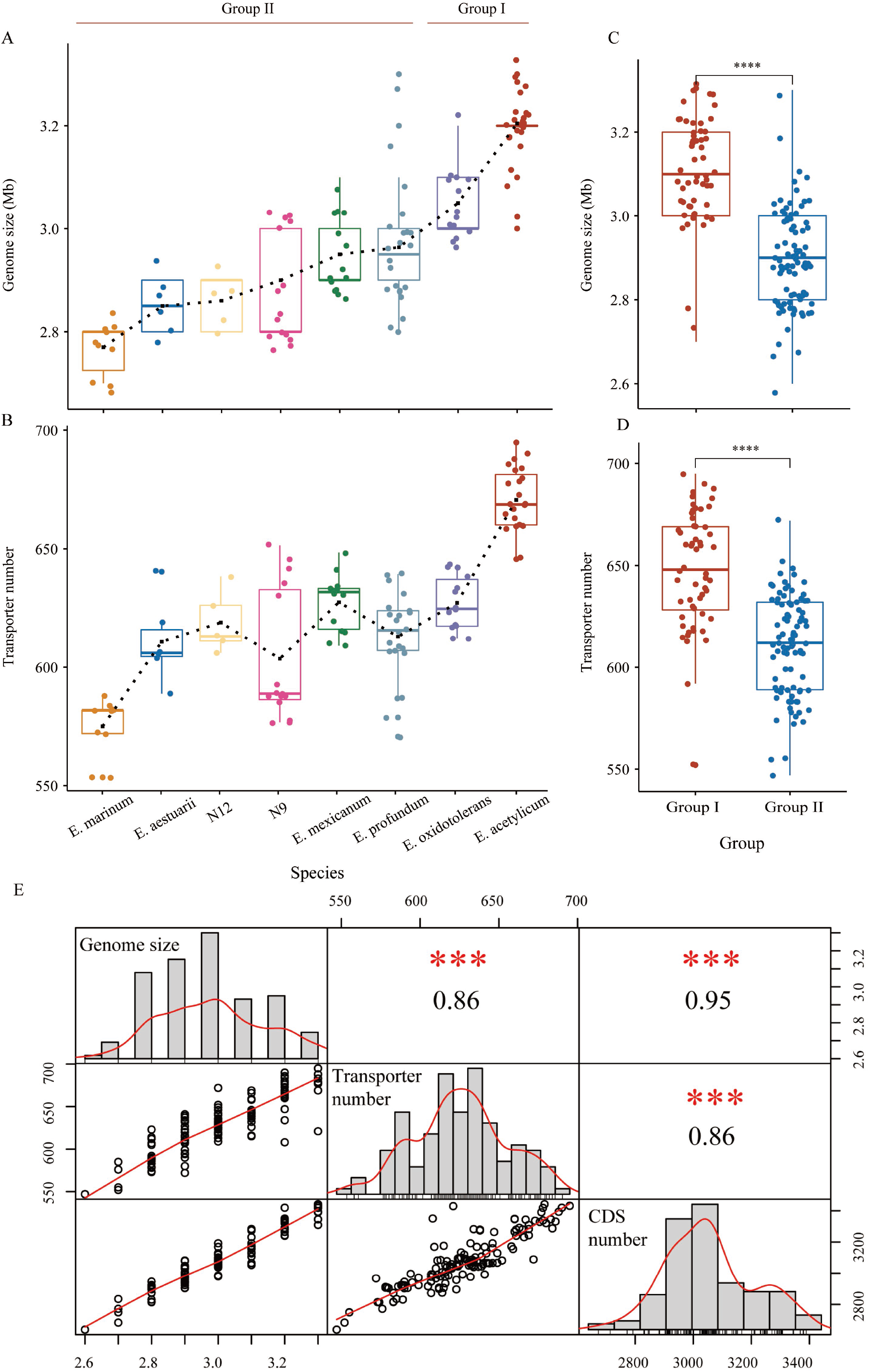
Comparison and Spearman’s correlation analysis of genome size with transporter. (A) Trends in genome size of *Exiguobacterium* species (the species with 5 strains were selected to show the trends). (B) Trends in transporter number of *Exiguobacterium* species (the species with 5 strains were selected to show the trends). (C) Comparison of genome size between Group I and Group II (the black ‘****’ represents significantly different, *p* < 0.0001, Wilcoxon tests). (D) Comparison of transporter number between Group I and Group II. (E) Spearman’s correlation analysis of genome size, CDS number and transporter number (the genome size, CDS number and transporter number is shown in the central diagonal; the scatterplots are depicted with a fitted red line and on the corresponding side for each pairing is the Spearman rank correlation coefficient *r_s_,* the red asterisks ‘***’ represents significance levels, *p* < 0.01).

Transporters are vital to all living organisms in the uptake of nutrients, secretion of metabolites, maintenance of ion concentration gradient across membranes and efflux of drug and toxins (50). In this study, 25 of the 247 identified transporter families were identified to have high degrees of correlation with both genome size and CDS number (correlation coefficients > 0.6, *p* < 0.01, Table 1). These 25 transporter gene families were significantly enriched in genome of Group I compared to Group II (Fig. 6; Table S9).

**Fig. 6.**
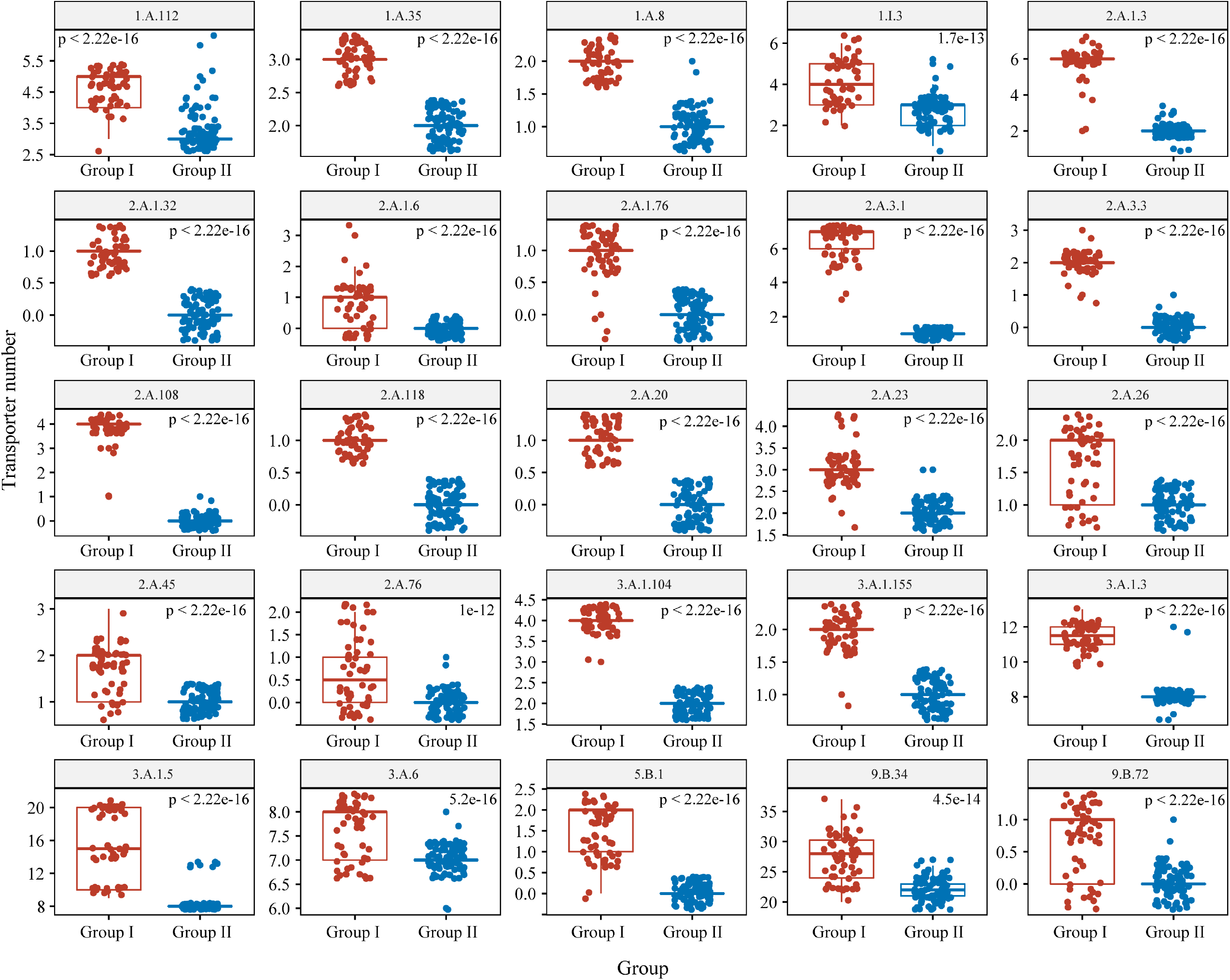
Comparison of 25 transporter family between Group I and Group II. All pairwise comparisons were significantly different (Wilcoxon test).

**Table 1.** Correlation analysis of transporter family with both genome size and CDS number.

| Genome content | TC number | Family | Correlation coefficients | p-value |
| --- | --- | --- | --- | --- |
| Genome size | 2.A.1 | **The Major Facilitator (MFS) Superfamily** | 0.73 | 0 |
| CDS number | 2.A.1 |  | 0.66 | 0 |
| Genome size | 2.A.1.3 | The Drug:H<sup>+</sup> Antiporter-2 (14 Spanner) (DHA2) Family | 0.73 | 0 |
| CDS number | 2.A.1.3 |  | 0.66 | 0 |
| Genome size | 2.A.1.76 | The Uncharacterized Major Facilitator 24 Family | 0.69 | 0 |
| CDS number | 2.A.1.76 |  | 0.61 | 4.44E-16 |
| Genome size | 2.A.1.32 | The Putative Aromatic Compound/Drug Exporter (ACDE) Family | 0.68 | 0 |
| CDS number | 2.A.1.32 |  | 0.60 | 8.88E-16 |
| Genome size | 2.A.1.6 | The Metabolite:H<sup>+</sup> Symporter (MHS) Family | 0.66 | 0 |
| CDS number | 2.A.1.6 |  | 0.67 | 0 |
| Genome size | **2.A.3** | **The Amino Acid-Polyamine-Organocation (APC) Superfamily** | 0.67 | 0 |
| CDS number | **2.A.3** |  | 0.61 | 2.22E-16 |
| Genome size | 2.A.3.1 | The Amino Acid Transporter (AAT) Family | 0.71 | 0 |
| CDS number | 2.A.3.1 |  | 0.66 | 0 |
| Genome size | 2.A.3.3 | The Cationic Amino Acid Transporter (CAT) Family | 0.68 | 0 |
| CDS number | 2.A.3.3 |  | 0.60 | 8.88E-16 |
| Genome size | **3.A.1** | **The ATP-binding Cassette (ABC) Superfamily** | 0.75 | 0 |
| CDS number | **3.A.1** |  | 0.69 | 0 |
| Genome size | 3.A.1.5 | The Peptide/Opine/Nickel Uptake Transporter (PepT) Family | 0.72 | 0 |
| CDS number | 3.A.1.5 |  | 0.68 | 0 |
| Genome size | 3.A.1.3 | The Polar Amino Acid Uptake Transporter (PAAT) Family | 0.73 | 0 |
| CDS number | 3.A.1.3 |  | 0.66 | 0 |
| Genome size | 3.A.1.155 | The Phage Infection Protein (PIP) Family | 0.72 | 0 |
| CDS number | 3.A.1.155 |  | 0.63 | 0 |
| Genome size | 3.A.1.104 | The Teichoic Acid Exporter (TAE) Family | 0.68 | 0 |
| CDS number | 3.A.1.104 |  | 0.60 | 1.33E-15 |
| Genome size | 1.A.112 | The Cyclin M Mg<sup>2+</sup> Exporter (CNNM) Family | 0.72 | 0 |
| CDS number | 1.A.112 |  | 0.70 | 0 |
| Genome size | 1.A.8 | The Major Intrinsic Protein (MIP) Family | 0.70 | 0 |
| CDS number | 1.A.8 |  | 0.61 | 2.22E-16 |
| Genome size | 1.A.35 | The CorA Metal Ion Transporter (MIT) Family | 0.68 | 0 |
| CDS number | 1.A.35 |  | 0.60 | 8.88E-16 |
| Genome size | 1.I.3 | The Bacterial (Planctomycetes) Nuclear Pore-like Complex (B-NPC) Family | 0.63 | 0 |
| CDS number | 1.I.3 |  | 0.62 | 2.22E-16 |
| Genome size | 2.A.23 | The Dicarboxylate/Amino Acid:Cation (Na<sup>+</sup> or H<sup>+</sup>) Symporter (DAACS) Family | 0.71 | 0 |
| CDS number | 2.A.23 |  | 0.65 | 0 |
| Genome size | 2.A.26 | The Branched Chain Amino Acid: Cation Symporter (LIVCS) Family | 0.61 | 4.44E-16 |
| CDS number | 2.A.26 |  | 0.61 | 2.22E-16 |
| Genome size | 2.A.45 | The Arsenite-Antimonite (ArsB) Efflux Family | 0.60 | 1.11E-15 |
| CDS number | 2.A.45 |  | 0.62 | 0 |
| Genome size | 2.A.118 | The Basic Amino Acid Antiporter (ArcD) Family | 0.68 | 0 |
| CDS number | 2.A.118 |  | 0.60 | 8.88E-16 |
| Genome size | 2.A.20 | The Inorganic Phosphate Transporter (PiT) Family | 0.68 | 0 |
| CDS number | 2.A.20 |  | 0.60 | 8.88E-16 |
| Genome size | 2.A.108 | The Iron/Lead Transporter (ILT) Family | 0.69 | 0 |
| CDS number | 2.A.108 |  | 0.61 | 4.44E-16 |
| Genome size | 2.A.76 | The Resistance to Homoserine/Threonine (RhtB) Family | 0.61 | 8.88E-16 |
| CDS number | 2.A.76 |  | 0.61 | 4.44E-16 |
| Genome size | 3.A.6 | The Type III (Virulence-related) Secretory Pathway (IIISP) Family | 0.62 | 0 |
| CDS number | 3.A.6 |  | 0.62 | 0 |
| Genome size | 5.B.1 | The gp91<sup>phox</sup> Phagocyte NADPH Oxidase-associated Cytochrome b<sub>558</sub> (Phox) Family | 0.73 | 0 |
| CDS number | 5.B.1 |  | 0.67 | 0 |
| Genome size | 9.B.34 | The Kinase/Phosphatase/Cyclic-GMP Synthase/Cyclic di-GMP Hydrolase (KPSH) Family | 0.75 | 0 |
| CDS number | 9.B.34 |  | 0.76 | 0 |
| Genome size | 9.B.72 | The 4 TMS GlpM (GlpM) Family | 0.63 | 0 |
| CDS number | 9.B.72 |  | 0.63 | 0 |

Seven of the 25 families are associated with the transport of diverse amino acids. Among them, cationic amino acid, polar amino acid, branched chain amino acid and basic amino acid are important components for nitrogen metabolism, protein synthesis, cell growth and energy production or conversion (51). Besides, 2 of the 25 families are involved in the transport of Mg^2+^, including the cyclin M Mg^2+^ exporter family and the CorA metal ion transporter family. Mg^2+^ homeostasis is important in bacteria and has been reported to play a critical role in their thermotolerance (52, 53). Moreover, the inorganic phosphate transporter family was also expanded in Group I. Compared with the marine environment, the terrestrial environment is more diverse and has a variety of complex microenvironments due to the influence of climate or seasons, which lead to a diverse content of nutrient substrates (3, 54). The enhanced capability of Group I strains in important substrates transport meets the need of cellular metabolism and functions, and provides important base to survive in more diverse terrestrial environments.

Three of the 25 transporter families are involved in transport of heavy metals ions, including the arsenite-antimonite efflux family, the iron/lead transporter family and peptide/opine/nickel uptake transporter family. These transporters have been shown to counteract the effects of toxic heavy metals (55–57). The members of major facilitator superfamily (MFS) are capable of transporting a wide range of substrates in response to ion gradients or function as drug:H^+^ antiporter; and majority of bacterial drug efflux pumps classified within the MFS (58, 59). Four of the 25 families are belonging to MFS superfamily, among which 2 are associated with drug efflux. Moreover, 2 of the 25 families are involved in the formation of bacterial cell wall and biofilm, including the teichoic acid exporter family and the 4 TMS GlpM family. Teichoic acid is a major cell wall component of Gram-positive bacteria, and has been proved to play crucial roles in bacterial resistance to antimicrobial and survival under disadvantageous conditions (60, 61). The members of 4 TMS GlpM family is required for normal production of alginate (62). Alginates are important polymeric substances contributing to the formation and development of biofilm matrixes of numerous bacteria enhancing colonization and persistence under environmental stresses (63). Due to the emission or leach out from the industrial and agricultural fields, terrestrial and freshwater ecosystems are contaminated with heavy metals or pesticide severely (64, 65). The expansion of transporters involved in disadvantageous conditions resistance and efflux of drug and heavy metals may contribute a more diverse distribution in non-marine habitats of Group I strains.

Efficient transport of substances related to metabolism, cellular function or environment stresses resistance is crucial for bacterial survival in a variety of environments (66). In order to survive in more diverse environments, bacteria have to develop specific systems for their survival such as nutrient sensing and transport systems (67). Bacteria are often exposed to stress conditions in many stages of their life cycle; and the capacity of environmental stress resistance determines the distribution of microorganisms (68, 69). Therefore, these expanded families that related to environment stresses resistance and transport of organic or inorganic substrates, may play crucial roles for *Exiguobacterium* spp. from Group I to survive in more diverse environments.

### Conclusions

Numerous studies have investigated the evolution and adaptation mechanism of important pathogens (70). However, fewer studies have addressed how microbes with cosmopolitan distribution but relative low abundance to adapt to diverse habitats. The wide distribution characteristic makes the genus *Exiguobacterium* as a valuable model for studying the adaptive strategy of bacteria to multiple habitats. This study suggested that these bacteria with nomadic lifestyle and cosmopolitan distribution are usual generalists, which can utilize a variety of nutrients simultaneously and resist diverse environmental stresses.

Although strains of *Exiguobacterium* genus are generalists, the ecological difference between members of Group I and II was still discovered. We found the species from Group I was distributed in more diverse environments with larger genome size and the expansion of transporter families contributed to the difference of genome size. Most of the enriched transporter families are involved in environment stresses resistance and transport of organic or inorganic substrates, which may play vital role for Group I strains in adaptation to wider habitats.

## MATERIALS AND METHODS

### Analysis of *Exiguobacterium* 16S rRNA gene sequences

We retrieved *Exiguobacterium* 16S rRNA gene sequences from GenBank. The information of isolation source of these sequences was collected. The habitats were classified into 13 types, including air, animal-associated environments, cold-associated environments, estuary-associated environments, contaminated water or soil, freshwater or freshwater sediment, hot-associated environments, hypersaline environments, marine-associated environments, plant, rhizosphere, soil and other inland environments.

### Bacterial isolation and culture

A total of 103 strains were used for adaptive experiments and genome sequencing, including 98 isolated from terrestrial and marine environments worldwide by us, and 5 type strains obtained from DSMZ and CGMCC (Table S2). Initially samples from marine and terrestrial environments were macerated and mixed with sterile saline solution (0.8%) using a standard dilution plating method on Marine agar 2216 (MA, Difco) and LB agar at 20 °C, respectively. All these 103strains can grow in sea salts free medium and were routinely cultivated on LB agar and in liquid LB for subsequent genomic sequencing.

### Adaptive ability tests of pH, temperature and salinity

To assess the range of adaptation to pH, temperature and salinity, all of the 103 strains were measured for assessing growth under the following conditions. The temperature range for growth were measured at 4, 25, 30, 40, 45 and 55 °C (the lowest growth temperature was not recorded for the strain that could not grow at 4°C) on LB and in liquid LB. The pH range for growth was determined at 25 °C in liquid LB medium at pH 4.0-12.0 (with intervals of 1 units) using the following buffers: citrate/Na_2_HPO_4_ buffer (pH 4.0-7.0), Tris buffer (pH 7.5-9.0) and NaHCO_3_/Na_2_CO_3_ buffer (pH 9.5-10.0); growth was evaluated by measuring OD600 after 7 days of incubation. Growth with 0—15% (in 1 % increments, w/v) NaCl was investigated after 14 days of cultivation at 25 °C in the following medium: 0.1% peptone, 0.1% yeast extract, 0.03% KCl, 0.25% MgSO_4_.7H_2_O, 0.05% CaCl_2_.

### Degradation ability tests of complex carbohydrates and proteins

Protease and amylase activities were tested on LB agar supplemented with skim milk (2%, w/v) and starch (0.4%, w/v). After 4—6 days at 25°C, a positive reaction was noticed when transparent zones around the colonies were directly visible or detected after coloration of the undegraded substrate.

### Genome sequencing, assembly and annotation

Genomic DNA was extracted by using a bacterial genomic DNA Mini kit (TaKaRa Bio) following the manufacturer’s protocol. Genomes of the 103 *Exiguobacterium* strains were sequenced using Illumina NovaSeq 6000 platform. The raw reads of each genome were trimmed using trimmomatic v0.36 (71) and *de novo* assembled using (25). The genomes of *Exiguobacterium* deposited in GenBank were collected and filtered based on the criterion that genomes were at least 95% complete, with < 5% contamination based on CheckM analysis (72). The ANI values between two each genome pair was computed using the OrthoANI software (73). Gene predictions and annotations of all genomes were generated using Prokka (74).

### Phylogenetic tree construction

Analysis of orthologous clusters was performed using the FastOrtho (http://enews.patricbrc.org/fastortho/), a faster reimplementation of OrthoMCL (75). In brief, an all-against-all BLAST was firstly performed with *E*-values < 1 × 10^−5^. Then, ortholog groups were created with the MCL algorithm with an inflation value of 2, and the single-copy gene families were obtained using custom-made Python scripts. Protein sequences of each family were aligned by MUSCLE (76) and then trimmed by trimAL (77). All trimmed alignments were then concatenated into a new alignment by a local Python script. Single-copy core gene-based phylogenetic tree was constructed using RAxML (78) with 1000 bootstrap replicates, employing the LG+I+G+F model. iTOL was used for the phylogenetic tree visualization (79).

### Identification of carbohydrate-active enzymes and proteases

For genes encoding carbohydrate-active enzymes and proteases, all of the annotation genes were searched against the CAZy database (www.cazy.org) (80) and peptidase database (MEROPS) (81) with *E*-values < 1 × 10^−5^ by BLASTP. The potential secreted carbohydrate-active enzymes and peptidases were confirmed based on the identification of extracellular transport signals using SignalP (82). Genes related to carbohydrate-active enzymes and proteases were further classified into different groups according to the predictions.

### Identification of vital genes for environmental stresses resistance

For each predicted protein by Prokka were annotated using BLASTP and Hmmscan against Clusters of Orthologous Groups (COG) database and PFAM database with *E*-values < 1 × 10^−5^, respectively. Transporters were predicted by performing BLASTP with *E*-values < 1 × 10^−10^ using *Exiguobacterium* protein sequences against all Transporter Classification Database (TCDB) sequences (83).

### Data analyses

Statistical analyses were performed using Wilcoxon test. Correlation analysis on the genome size, CDS number and transporter number was performed using chart.Correlation from the PerformanceAnalytics package in R (https://cran.r-project.org/web/packages/PerformanceAnalytics/index.html).

### Data availability

The genomes supporting the results have been deposited at DDBJ/ENA/GenBank under the BioProjectID PRJNA644789 (accession numbers from JACSJK000000000 to JACSNI000000000) (Table S2).

## ACKNOWLEDGEMENTS

This study was supported by National Natural Science Foundation of China (31670002, 31970003 and 31770003).

We declare that we have no conflict of interest.

---

**Fig. S1 Validation of the ability of *Exiguobacterium* spp. to degrade and metabolize complex carbohydrates and peptides.**

**Table S1 Information of *Exiguobacterium* 16S rRNA gene sequences in Genebank.**

**Table S2 Genome features and isolation source of 145 *Exiguobacterium* strains.**

**Table S3 Temperature, pH and salinity tolerance test of 103 *Exiguobacterium* strains.**

**Table S4 Average nucleotide identity (ANI) analysis of 145 *Exiguobacterium* strains.**

**Table S5 Niches distribution of 145 *Exiguobacterium* strains basing on phylogenetic.**

**Table S6 Gene number of carbohydrate-active enzymes (CAZymes) detected in each *Exiguobacterium* genome.**

**Table S7 Gene number of extracellular peptidases detected in each *Exiguobacterium* genome using the MEROPS peptidase database.**

**Table S8 Genetic basis for *Exiguobacterium* strains adaptation to diverse habitats.**

**Table S9 Transporter analysis of 145 *Exiguobacterium* genomes.**

